# XWAS: a software toolset for genetic data analysis and association studies of the X chromosome

**DOI:** 10.1101/009795

**Authors:** Feng Gao, Diana Chang, Arjun Biddanda, Li Ma, Yingjie Guo, Zilu Zhou, Alon Keinan

**Affiliations:** Department of Biological Statistics and Computational Biology, Cornell University, Ithaca, NY 14853, USA; Program in Computational Biology and Medicine, Cornell University, Ithaca, NY 14853, USA; Department of Animal and Avian Sciences, University of Maryland, College Park, MD 20740, USA; School of Computer Science and Technology, Harbin Institute of Technology, Harbin, Heilongjiang 150001, China

**Keywords:** GWAS, software, X chromosome, genetic association study, complex diseases

## Abstract

XWAS is a new software suite for the analysis of the X chromosome in association studies and similar studies. The X chromosome plays an important role in human disease, especially those with sexually dimorphic characteristics. Special attention needs to be given to its analysis due to the unique inheritance pattern, which leads to analytical complications that have resulted in the majority of genome-wide association studies (GWAS) either not considering X or mishandling it with toolsets that had been designed for non-sex chromosomes. We hence developed XWAS to fill the need for tools that are specially designed for analysis of X. Following extensive, stringent, and X-specific quality control, XWAS offers an array of statistical tests of association, including: (1) the standard test between a SNP (single nucleotide polymorphism) and disease risk, including after first stratifying individuals by sex, (2) a test for a differential effect of a SNP on disease between males and females, (3) motivated by X-inactivation, a test for higher variance of a trait in heterozygous females as compared to homozygous females, and (4) for all tests, a version that allows for combining evidence from all SNPs across a gene. We applied the toolset analysis pipeline to 16 GWAS datasets of immune-related disorders and 7 risk factors of coronary artery disease, and discovered several new X-linked genetic associations. XWAS will provide the tools and incentive for others to incorporate the X chromosome into GWAS, hence enabling discoveries of novel loci implicated in many diseases and in their sexual dimorphism.

## Introduction

Genome-wide association studies (GWAS) have identified thousands of loci underlying complex human diseases and other complex traits (Welter, *et al.*, 2014). While successful for the autosomes (non-sex chromosomes), the vast majority of these studies have either incorrectly analyzed or ignored the X chromosome (X) (Wise, Gyi and Manolio, 2013). In most studies, all variants on the X chromosomes have been removed as a consequence of the quality control (QC) procedures (Mailman, *et al.*, 2007; Wise, *et al.*, 2013; Chang, *et al.*, 2014; Tryka, *et al.*, 2014). Many other studies that did analyze the X chromosome incorrectly applied methods that have been designed for the autosomes, without accounting for the analytical problems arising from X’s unique mode of inheritance and its consequent population genetic and evolutionary patterns (Hammer, *et al.*, 2008; Wilson and Makova, 2009; Emery, Felsenstein and Akey, 2010; Hammer, *et al.*, 2010; Keinan and Reich, 2010; Lambert, *et al.*, 2010; Lohmueller, Degenhardt and Keinan, 2010; Arbiza, *et al.*, 2014). As a result, the role X plays in complex diseases and traits remains largely unknown.

Many human diseases commonly studied in GWAS show sexual dimorphism, including autoimmune diseases (Voskuhl, 2011), cardiovascular diseases (Lerner and Kannel, 1986) and cancer (Muscat, *et al.*, 1996; Matanoski, *et al.*, 2006), which suggests a potential contribution of the X chromosome (Carrel and Willard, 2005; Ober, Loisel and Gilad, 2008). Several recent studies have examined this issue and demonstrated the potential value of analyzing X (Chang, *et al*., 2014; Gilks, Abbott and Morrow, 2014; Tukiainen, *et al*., 2014; Ma, Hoffman and Keinan, 2015; Li YR, unpublished). However, while association methods, QC and analysis pipelines are well established for the autosomes, respective pipelines for X-linked data are not readily available. Hence, in this paper, we introduce the software package XWAS (chromosome X-Wide Analysis toolSet), which is tailored for analysis of genetic variation on X. It implements extensive functionality that carries out QC specially-designed for the X chromosome, statistical tests of single-marker association that account for its unique mode of inheritance, gene-based tests of association, and additional distinct tests only applicable to X that capitalize on its mode of inheritance. In implementing these features, the toolset builds on--and complements--the commonly-used PLINK (Purcell, *et al.*, 2007) software. It includes many novel features that can facilitate X-wide association studies that are not available in PLINK and, to the best of our knowledge, in any other software. Combined, the XWAS toolset integrates the X chromosome into GWAS as well as into the next generation of sequence-based association studies. (Chang, *et al.*, 2014; Gilks, Abbott and Morrow, 2014; Tukiainen, *et al.*, 2014; Ma, Hoffman and Keinan, 2015)

## Features and Functionality

### Quality Control Procedures

The XWAS toolset implements a whole pipeline for performing QC on genotype data for the X chromosome. The pipeline first follows standard GWAS QC steps as implemented in PLINK (Purcell, *et al.*, 2007) and SMARTPCA (Price, *et al.*, 2006) by running these tools. These include the removal of both individual samples and SNPs (single nucleotide polymorphisms) according to multiple criteria. Specifically, samples are removed based on (i) relatedness, (ii) high genotype missingness rate, and (iii) genetic ancestry differing from the majority of the samples (Price, *et al.*, 2006). SNPs are removed based on criteria such as their missingness rate, their minor allele frequency (MAF), and deviation from Hardy-Weinberg Equilibrium (HWE). While the toolset is currently focused on case-control GWAS (binary traits), the entire QC pipeline is also applicable to GWAS of quantitative traits. One filter applied only to binary traits is the removal of SNPs for which missingness is correlated with the trait, i.e. with case or control status (--*test-missing*).

To consider differences in genotyping between hemizygous males and diploid females, XWAS applies all the aforementioned QC steps of samples separately for males and females. Consequently, a unified dataset is generated for subsequent analyses that include all SNPs and individuals passing the above filtering criteria in both the male and female QC groups.

The pipeline then applies X-specific QC steps, which are exclusively built into XWAS, to the unified dataset. These include (i) removing SNPs with significantly different MAF between male and female samples in the control group (--*freqdiff-x*), (ii) removing SNPs with significantly different missingness rates between male and female controls (--*missdiff-x*), and (iii) the removal of SNPs in the pseudoautosomal regions (PARs). The first two of these steps capture problems in genotype calling when plates include both males and females (Korn, *et al.*, 2008). Further details regarding specific QC procedures can be found in the user manual that is available with the toolset.

### Single-Marker Association Testing on the X chromosome

For an X-linked SNP, while females have 0, 1, or 2 copies of an allele, hemizygous males have at most one copy. Via the process of X-inactivation, one of the two copies in females is usually transcriptionally silenced. If X-inactivation is complete, it produces monoallelic expression of X-linked protein-coding genes in females. Therefore, when considering loci that undergo complete X-inactivation, it may be apt to consider males as having 0/2 alleles, corresponding to the female homozygotes (the FM_02_ test). The toolset carries out this test for association between a SNP and disease risk by using the --*xchr-model 2* option in PLINK (Purcell, *et al.*, 2007). For other scenarios though, including where some genes on the X escape X-inactivation or different genes are inactivated in different cells, it can be more indicative to code males as having 0/1 alleles. Hence, the toolset further carries out such an association test (FM_01_ test) of a SNP by using the following options in PLINK (Purcell, *et al.*, 2007): *--logistic* and *--linear* for binary and quantitative traits, respectively.

All tests, including tests described in following sections, allow for covariates such as population structure, sex, and traits that are correlated with the disease, as commonly considered in GWAS. We suggest calculating principal components by using EIGENSTRAT (Price, *et al.*, 2006) and include them as covariates to control for population structure. Ten such principal components are considered by default, unless otherwise specified. Any other user-defined covariates can also be incorporated.

### Single-Marker Sex-stratified Analysis on the X chromosome

The XWAS software further includes new tests that are not included in PLINK. First, we implemented a new sex-stratified test, FM_comb_, which is particularly relevant for X analyses since SNPs and loci on the sex chromosomes are potentially more likely to exhibit different effects on disease risk between males and females. In such scenarios, as well as in scenarios where the effect is only observed in one sex, a sex-stratified test as described in the following can be better powered. This functionality is accessible by the option --*strat-sex*. The FM_comb_ test first carries out an association test separately in males and females and then combines the results of the two tests to obtain a final sex-stratified significance level. The combination of the two test statistics is implemented using both Fisher’s method (*--fishers*) (Fisher, 1925) (in the FM_F.comb_ test) and Stouffer’s method (*--stouffers*) (Stouffer, 1949) (in the FM_S.comb_ test).

Each of these two tests is more powerful in different scenarios (Chang, *et al.*, 2014), e.g. FM_F.comb_ allows the SNP tested to have different, even an opposite, effect on disease risk in males and females. FM_F.comb_ is also insensitive to whether males are coded as 0/2 (as in the FM_02_ test) or as 0/1 (as in the FM_01_ test), thus making no assumptions regarding X-inactivation status. Alternatively, FM_S.comb_ directly accounts for the potentially differing sample sizes between males and females to maximize power. For this latter test, XWAS weighs by the sample size in males and females in cases and controls following the approach of Willer *et al*. (2010). (Willer, Li and Abecasis, 2010)

### Single-Marker Sex-differentiated Effect Size Test on the X chromosome

We described above sex-stratified tests that accommodate associations with different effect size between males in females. In another type of test (FM_diff_), we directly test whether the effect size is different between the sexes. This test, applied to each SNP, runs a *t*-test to test for difference between the odds ratio (OR) in males alone and the OR in females alone, while accounting for hemizygosity in males. This test is implemented under the *--sex-diff* option and is further described in Chang *et al*. (2014). For this test and the sex-stratified test introduced in the previous section, both odds ratios and regression coefficients in each sex can be provided as output for further examination.

### Single-Marker Variance-based Testing Informed by X-inactivation in Females

During X-inactivation, the expression of one copy of the X chromosome in females is randomly silenced, thereby increasing variation in the expression of X-linked quantitative trait loci (QTL). Specifically, females that are heterozygotes for a QTL might exhibit higher phenotypic variance than homozygous females since one or the other allele might be more dominantly affecting the phenotype in each given female heterozygote, such that for some individuals the QTL expression is more similar to one type of female homozygous, while to the other type in other individuals. We developed a test aimed at capturing this increased variance as a means for detecting X-linked QTLs in females. An overview of the test and its implementation follows, while we refer readers to Ma *et al*. (2015) for a full description of the test. This test (*F*var) is currently implemented under the *--var-het* option. Although this *F*var test is implemented for quantitative traits, it can be generalized to qualitative traits by applying liability threshold modeling (Zaitlen, *et al.*, 2012) to transform disease status to an unobserved continuous liability.

The null hypothesis of the *F*_var_ test is that phenotypic variances of the three genotypic groups of a SNP with 0, 1, or 2 copies of a reference allele are all equal. The alternative hypothesis is that female heterozygotes show a higher phenotypic variance than others. Hypothesis testing is carried out using a modified Brown-Forsythe test of variances (Brown and Forsythe, 1974). We first normalize the phenotypic value and remove the effects of possible covariates by a linear regression as conventionally done, namely y = µ + XB + e, where *y* is a vector of quantitative trait levels, µ is the population mean, *X* is the matrix of possible covariates, and e is a vector of residuals. Assume *y*_*i|g=j*_ is the phenotypic value of the *i*^th^ individual in the *j*^th^ genotypic group and *z*_*i|g=j*_ = |*e*_*i|g=j*_| is the absolute residual value of the *i*^th^ individual in the *j*^th^ genotypic group (*j* = 0, 1, or 2 copies of an allele of a SNP). A test statistic is derived as

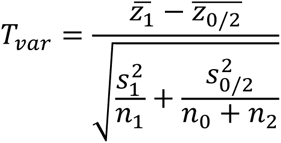

where 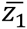 is the sample mean of *z*_*i|g=*1_ over *i*, 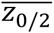 is the sample mean of *z*_*i|g=*0_ and *z*_*i|g=*2_ combined, 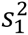 and 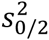 are the sample variances respectively, and *n*_*j*_ is the sample size of the *j*^th^ genotypic group. Under the null hypothesis, the statistic follows a *t*-distribution with degrees of freedom given by 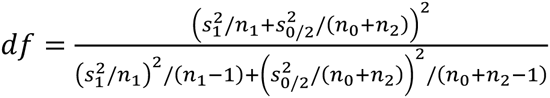.

This variance-based test captures a novel signal of X-linked associations by directly testing for higher phenotypic variance in heterozygous females than homozygotes. As a test of variance it is generally less powerful than standard tests of association that consider means; however, it provides an independent and complementary test to the standard association test for QTLs on X (Ma, *et al.*, 2015).

### X-linked Gene-based Testing

XWAS also includes unique features for carrying out gene-based association analysis on the X chromosome. Gene-based approaches may be better powered to discover associations than single-marker analyses in cases of a gene with multiple causal variants of small effect size, or of multiple markers that are each in incomplete linkage disequilibrium with underlying causal variant/s. Furthermore, in studying the effect of X on sexual dimorphism in complex disease susceptibility, it may be desirable to analyze whole-genes or all genes of a certain function combined based on their unique function or putatively differential effect between males and females, as illustrated in Chang *et al*. (2014).

The XWAS package determines the significance of association between each gene as a whole and disease risk by implementing a gene-level test statistic that combines individual SNP-level test statistics for all SNPs in and around each studied gene. This gene-level approach is applicable to any of the different tests described above. For instance, beyond tests of association, it can be applied to the sex-differentiated tests. In this case the gene-based test captures any scenario whereby SNPs within the gene display different effects in males and females, without restricting such differential effects to be of a similar nature across SNPs. By default, genes are considered from the UCSC browser “knownCanonical” transcript ID track. SNPs are mapped to a gene if they are in the gene or within 15 kb of the gene’s start or end positions. The user can also provide a different set of gene definitions or alternate regions of interest and a different window length around them in which SNPs are also to be considered.

Combining SNP statistics across a gene is implemented in the general framework of Liu *et al*. (2010). Specifically, the significance of an observed gene-based test statistic is assessed from the distribution of test statistics that is expected given the linkage disequilibrium between the SNPs in the gene. In Liu *et al*. (2010), the test statistics for all SNPs in the gene are summed. Here, we have implemented a slight modification to this procedure, whereby we combined SNP-based *p*- values with either the truncated tail strength (Jiang, *et al.*, 2011) or the truncated product (Zaykin, *et al.*, 2002) method, which have been suggested to be more powerful in some scenarios (Zaykin, *et al.*, 2002; Ma, Clark and Keinan, 2013).

To determine significance, XWAS follows the procedure in Liu *et al*. (2010). The observed statistic is compared to gene-level test statistics obtained when SNP-level statistics are randomly drawn from a multivariate normal distribution with a covariance determined by the empirical linkage disequilibrium between the SNPs in the tested gene. The significance level is then the proportion, out of many such drawings, for which this sampled gene-level statistic is more, or as extreme compared to the empirical one. For computational efficiency, the number of drawings is determined adaptively as in Liu *et al*. (2010). By combining the truncated tail measures with this procedure, our new gene-based method combines the test statistics from multiple SNPs that show relatively low *p*-values, while also accounting for the dependency between these *p*-values due to linkage disequilibrium between the SNPs. Such a *p*-value is estimated for each gene and for each of the X-linked tests described above. (Liu, *et al.*, 2010)

## Examples of Use

In this section, we summarize several sets of results obtained using the XWAS software and publicly available GWAS datasets. For many of the results, we include herein a brief description of the main results, with the full description appearing in separate papers (Chang, *et al.*, 2014; Ma, *et al.*, 2015). All associations presented herein are significant and details regarding the *p*- values can be found in the respective papers.

### Association of X-linked SNPs with Autoimmune Diseases

We applied the XWAS software described above to 16 GWAS datasets of autoimmune disease and other disorders with a potential autoimmune-related component. These include the following datasets that we obtained from dbGaP (Mailman, *et al.*, 2007; Tryka, *et al.*, 2014): ALS Finland (Laaksovirta, *et al.*, 2010) (phs000344), ALS Irish (Cronin, *et al.*, 2008) (phs000127), Celiac disease CIDR (Ahn, *et al.*, 2012) (phs000274), MS Case Control (Baranzini, *et al.*, 2009) (phs000171), Vitiligo GWAS1 (Jin, *et al.*, 2010) (phs000224), CD NIDDK (Duerr, *et al.*, 2006) (phs000130), CASP (Nair, *et al.*, 2009) (phs000019), and T2D GENEVA (Qi, *et al.*, 2010) (phs000091). Similarly, we obtained the following datasets from the Wellcome Trust Case Control Consortium (WT): all WT1 (The Wellcome Trust Case Control Consortium, 2007) datasets, WT2 ankylosing spondylitis (Evans, *et al.*, 2011), WT2 ulcerative colitis (UK IBD Genetics Consortium, *et al.*, 2009) and WT2 multiple sclerosis (International Multiple Sclerosis Genetics Consortium, *et al.*, 2011). Finally, we also analyzed data from Vitiligo GWAS2 (Jin, *et al.*, 2012). These datasets are described in more detail in Chang *et al*. (2014).

Following application of the QC pipeline as described above, we applied the SNP-level FM_02,_ FM_F.comb_, and FM_S.comb_ tests to all SNPs in each of the 16 datasets. Based on the Vitiligo GWAS1 datasets, we associated SNPs in a region 17 kilobases (kb) away from the retrotransposed gene retro-*HSPA8* with risk of vitiligo. The parent of this retrotransposed gene, *HSPA8* on chromosome 11, encodes a member of the heat shock protein family, which has been previously associated to vitiligo (Mosenson, *et al.*, 2012; Abdou, Maraee and Reyad, 2013; Mosenson, *et al.*, 2013). We discovered another association in WT2 ulcerative colitis of SNPs in an intron of *BCOR* contributing to ulcerative colitis disease risk. BCOR indirectly mediates apoptosis via co-repression of *BCL-6* (Huynh, et al., 2000).

### Association of Whole X-linked Genes with Autoimmune Diseases

We next focused on a gene-based analysis of the X chromosome by using the SNP-level results of all the three tests in the above results as a basis for gene-based tests in the same 16 datasets. This analysis led to the discovery of the first X-linked gene-based associations with any disease or trait, which supports the utility of the XWAS package in facilitating such analyses. We associated in Vitiligo GWAS1 and replicated in Vitiligo GWAS2 an association between the gene *FOXP3* and vitiligo disease risk, in support of an earlier candidate gene study (Birlea, *et al.*, 2011). We also found a novel association of *ARHGEF6* to Crohn’s disease and further replicated it in ulcerative colitis, another inflammatory bowel disorder (IBD). ARHGEF6 binds to a surface protein of a gastric bacterium (*Helicobacter pylori*) that has been associated to IBD (Luther, *et al.*, 2010; Jin, *et al.*, 2013). Finally, we associated *CENPI* as contributing to the risk of three different diseases (amyotrophic lateral sclerosis, celiac disease and vitiligo). Other, autosomal genes in the same family as *CENPI* have previously been associated to amyotrophic lateral sclerosis (Ahmeti, *et al.*, 2013) as well as multiple sclerosis (Baranzini, *et al.*, 2009), supporting an involvement of *CENPI* with autoimmunity in general.

### X-linked SNPs Showing Sex-differentiated Effect Size with Autoimmune Disease

As a final analysis on the 16 autoimmune datasets, we applied the FM_diff_ test and its gene-based version. Based on this test, we discovered and replicated the gene *C1GALT1C1* (also known as *Cosmc*) as exhibiting sex-differentiated effect size in risk of IBD. *C1GALT1C1* is necessary for the synthesis of many O-glycan proteins (Ju and Cummings, 2005), which are components of antigens. We further found *CENPI*, which we previously associated with several diseases, to show significantly different effects in males and females in the same diseases as in the association analysis.

### Increased Variance of Systolic Blood Pressure in Heterozygous Females for an X-linked SNP

As an example application of the variance-based testing informed by X-inactivation, we considered data on 7 quantitative traits from the Atherosclerosis Risk in Communities (ARIC) study (Williams, 1989) along with Affymetrix 6.0 data from the participating individuals, which included 34,527 X-linked SNPs. First, we applied the entire set of QC procedures implemented in XWAS for quantitative traits. Then, we applied our single-marker variance-based testing and compared with application of standard testing for a QTL. Across the 7 traits, we found one SNP with a significant association based on the variance test (Ma, *et al.*, 2015). Importantly, the signals of this test are not in the same loci as those of the standard test, in line with them capturing different types of signals. Specifically, the significant SNP, rs4427330, which is associated with systolic blood pressure based on the variance test, is not associated with any trait based on the standard test. It is located upstream of *AFF2*, which might regulate *ATRX*. *ATRX* is associated with alpha-thalassemia, a disease that can cause anemia and has been associated with hypertension (Bowie, Reddy and Beck, 1997).

## Implementation and Availability

The XWAS software package is implemented in C++ and includes in part functions from open-source PLINK (Purcell, *et al.*, 2007). This software uses the same input format as PLINK. Beyond C++, additional features are also implemented in scripts, including in shell (for QC), Perl (for converting file formats and using SMARTPCA), and R (for gene-based testing). The entire package is freely available for download from http://keinanlab.cb.bscb.cornell.edu and includes (1) scripts, (2) the binary executable XWAS, (3) all source code with a Makefile, (4) a user manual, and (5) example data and examples of running the different options offered by the package. Additional help is provided via the --*xhelp* option. The XWAS toolset was initially designed and optimized for Linux systems, hence exhibits best performance in such systems. A Makefile is also provided to facilitate local compilation on Linux environments, and can also be adjusted for Windows and MAC OS by revising a few lines indicated therein.

## Conclusions

We have developed XWAS, an extensive toolset that facilitates the inclusion of the X chromosome in genome-wide association studies. It offers X-specific QC procedures, a variety of X-adapted tests of association, and an X-specific test of variance testing, available for both single-marker and gene-based statistics. We applied this toolset to successfully discover and replicate a number of genes with autoimmune disease risk and blood pressure.

We are continually developing the software and upcoming versions in the near future will offer additional features, including all features needed to conduct an extensive association study of quantitative traits (many features for quantitative traits are already functional in the current version). Similarly, while imputation of unobserved SNPs is presently performed as a preprocessing step using IMPUTE2 (Howie, *et al.*, 2012), we will incorporate X-specific imputation as part of the pipeline. Additional features will include analysis of X-linked data from sequence-based association studies (including burden tests), statistical methods that have been previously designed for the X chromosome (Zheng, *et al.*, 2007; Clayton, 2008; Clayton, 2009; Loley, Ziegler and Konig, 2011; Thornton, *et al.*, 2012), additional tests we previously proposed based on the workings of X-inactivation (Ma, *et al.*, 2015), and tests for gene-gene interactions. Finally, we will incorporate information regarding whether or how often a gene undergoes or escapes X-inactivation (Carrel and Willard, 2005; Cotton, *et al*., 2011; Disteche, 2012; Slavney A, unpublished). For computational efficiency, we will also continually upgrade the functions of PLINK that XWAS uses to the most recent version. (Carrel and Willard, 2005; Cotton, *et al.*, 2011; Disteche, 2012)

This software, and through incorporation of additional features, can be used for other types of studies of the X chromosome beyond association studies, in particular population genetic studies. For instance, allele frequency output and testing for significant differences in allele frequency between males and females as currently implemented, can be utilized to search for signals of selection.

Considering the availability of unutilized data for the X chromosome from thousands of GWAS, and the additional X-linked data that is being generated as a part of ongoing GWAS, many researchers will find extensive utility in the XWAS toolset. Furthermore, it is not limited to application to human data, but rather genetic data from all organisms with XX/XY sex determination system, including all mammals. XWAS will facilitate the proper analysis of these data, incorporate X into GWAS and enable discoveries of novel X-linked loci as implicated in many diseases and in their sexual dimorphism.

## Acknowledgements

We thank Paul Billing-Ross, Aviv Madar, Aaron Sams, Andrea Slavney, Richard Spritz, Yedael Y. Waldman and Liang Zhang for helpful comments on the software and previous versions of this manuscript.

